# Global convergence in wood evolution is driven by drought on continents and frost-free temperatures on islands

**DOI:** 10.1101/2025.09.15.676260

**Authors:** Alexander Zizka, Matthias Grenié, Timo Conradi, Renske E. Onstein, Helge Bruelheide, Frederic Lens

## Abstract

Phylogenetically derived woodiness (DW), the evolutionary reversion from herbaceousness to woodiness in angiosperms, is one of the most conspicuous characteristics of (sub)tropical island floras. Here, we show that DW across continents is more common than previously thought, especially in frost-free and open habitats with pronounced seasonal drought. Using a novel dataset on the evolution of woodiness in angiosperms, we discovered substantially more derived woody species (DWS) and independent evolutionary transitions on continents compared to islands (4,808 species and 513 transitions vs. 1,084 and 175, respectively). However, we identified more insular DWS hotspots (22) than the four continental DWS hotspots: the Andes, Southern Africa, the Old-World Dry Belt and Australia. A structural equation model controlling for total species richness suggests that aridity is the strongest predictor of the number of DWS on continents, while frost-free temperatures best predict DWS on islands. Precipitation seasonality and mean elevation emerge as additional significant predictors in both cases, with a further potential role for light-prone open habitats. In summary, the diverse global drivers behind the hundreds of independent woodiness shifts highlight the existence of various mechanisms that lead to increased wood formation in stems, confirming its adaptive value over evolutionary time.

## Introduction

The evolutionary history of wood, a lignin-rich plant tissue produced by the vascular cambium ^1^, goes back more than 400 Ma ^2,3^. The paleoclimatic conditions in which the first woody species emerged remain unknown, making it difficult to formulate well-founded hypotheses about why early land plants evolved woody growth. Hence, repeated transitions towards wood formation in herbaceous lineages in more recent times offer insights into the processes underlying woodiness evolution. Indeed, biologists have been fascinated for centuries by the convergent evolution of woodiness in herbaceous angiosperm lineages after colonising (sub)tropical islands that often resulted in iconic woody island radiations (insular woodiness, IW) ^4–8^.

As DW is mostly described from (sub)tropical islands, most hypotheses on its drivers rely on island conditions. Primarily, IW has been linked to a longer plant life span either resulting from the surrounding ocean causing a more stable favourable frost-free climate ^4^, or from the absence of large mammal herbivores that would otherwise eat small fragile plants ^4^, or from pollinator scarcity favouring plants with a longer flowering time required for cross-pollination in the small population(s) ^8^. Alternatively, herbaceous colonisers in the open environments of newly emerged islands may be selected to grow taller and hence develop mechanically stronger woody stems, either to gain a competitive advantage for light ^5,9^ or to avoid burial under volcanic deposits ^10^. An alternative hypothesis is that drought is a major driver of DW. This is supported by multiple lines of evidence from the Canary Islands: (1) the distribution of insular woody species (IWS), which predominantly thrive in drier vegetation types ^7^, (2) the timing of island colonization of most IW lineages during dry paleoclimatic periods ^11^, and (3) the positive role of increased stem woodiness in delaying lethal levels of root-to-leaf water transport blockage by drought-induced gas bubbles in the xylem vessels ^12^. Moreover, drought (and lack of herbivory) have been identified as global predictors of IWS richness across oceanic and continental islands ^13^.

Several lines of evidence suggest that DW may be more common on continents than currently assumed in the literature. First, 800 island species evolved their woodiness on continents before colonising islands ^13^. Second, an increasing number of well-resolved phylogenies reveals that DW has re-evolved repeatedly in distantly related lineages across various habitats on continents ^14–20^. Particularly iconic radiations are known from mountains, such as the giant groundsels (*Dendrosenecio* spp.) in East Africa and the rosette tree species of *Espeletia* in the South American paramo ^21,22^. Third, most herbaceous angiosperm species (excluding monocots) keep the ability to develop a vascular cambium that produces a limited amount of wood at the stem base ^15,18,23^. Fourth, molecular studies suggest that one or a few genes can turn the tiny herbaceous model plant *Arabidopsis thaliana* into a woody shrub phenotype, suggesting that the gene regulatory mechanism triggering wood formation could be activated easily on islands and continents ^24–27^. If continental DWS are indeed far more widespread than currently documented in the literature, this raises the question of whether the prevailing island-based hypotheses for IW can also account for the occurrence of DWS on continents.

To understand woodiness evolution across the globe, we expanded our previous database on IWS^13^ with a global literature review across angiosperms to also include continental DWS. This dataset provides a baseline for summarizing patterns of woodiness evolution and for evaluating the relevance of IW drivers in a continental context. Specifically, we test three hypotheses:

1. *The numbers of DWS and transitions to DW on continents are at least as high as on islands, but likely older due to the average younger island ages. Furthermore, continental DWS are geographically widespread and occur in the same families as IWS*.
2. *DW on continents and islands is driven by similar climatic factors. More specifically, we expect that the occurrence of DWS is positively related to frost-free conditions with recurrent drought cycles. Additional potential woodiness drivers may play a role on islands and/or continents*.
3. *The occurrence of DWS is higher in open habitats with little competition from tall woody trees, and lower in the dark understory of dense forests across the world*.

## Methods

### Data compilation

To identify DWS on continents, we compiled phylogenetically-inferred evidence on woodiness evolution from hundreds of floras, taxonomic revisions, herbarium specimens and stem cross sections in groups for which information on phylogenetic relationships were available (a total of 1,367 publications are included as references). To assess the evolutionary derived status of woodiness, we visually traced character evolution on these published phylogenies following a maximum parsimony approach to tease apart IWS (i.e., species evolved their woodiness on the islands after the arrival of an herbaceous continental coloniser), continental DWS (referring to species that underwent the transition from herbaceousness towards woodiness on the continent), and ancestrally woody species (i.e., species from a woody clade with only woody ancestors). We relied on this visual expert-based approach rather than modelled ancestral state reconstructions based on a supertree or supermatrix phylogeny of angiosperms, because the majority of DWS and their herbaceous close relatives are not included in available large-scale phylogenies ^28,29^ therefore biasing model reconstructions and making a global assessment of the biogeography and evolution of DW unreliable ^13^. Apart from identifying the woody species with herbaceous ancestry across flowering plants, we also summarized the number of individual evolutionary transitions identified per family and genus in the same way. Our approach enabled us to ensure a conservative estimate of the global number of DWS and the number of transitions to DW. Our estimates represent minimum numbers of DWS and DW transitions due to two reasons: (1) several phylogenies consulted did not show sufficient resolution and/or sampling to accurately address the origin of all the woody species within a given clade, and were therefore left out, and (2) we applied stringent criteria to define a woody species, thereby ignoring species with an intermediate herbaceous-woody habit.

Wood is defined as secondary xylem produced by a vascular cambium ^1^ In flowering plants that can produce wood, i.e. non-monocot angiosperms, the boundary between woody and herbaceous species is sometimes hard to define because of the continuous variation in wood development in the aboveground stem among woody and herbaceous species ^18,25^. Here, we only accepted a species to be “derived woody” under two criteria: (1) the species produces a distinct wood cylinder extending also towards the upper stem parts (i.e. shrubs, trees, lianas), thereby excluding species with only a woody stem base (“woody herbs” or suffrutescent species that are not woody enough according to our definition), and (2) the transition from herbaceousness towards derived woodiness can be clearly inferred from the available phylogenetic literature. The reason for ignoring the many species with an intermediate herbaceous-woody growth form is that we are only interested in disentangling the evolutionary processes that underlie the dramatic transitions from herbaceous species with no or limited wood formation in their basal stem parts to DWS with extensive wood development along substantial parts in their aboveground stems.

To separate DWS that evolved woodiness on continents (DWS on continents) from DWS that evolved woodiness on islands (IWS) versus DWS that evolved woodiness on the continent and subsequently colonized islands (DWS on islands), we recorded the geographic distribution of DWS and their relatives, and followed the same procedure using published phylogenies described above to identify if the woodiness evolution developed on an island or continent. Furthermore, we tried to obtain for each DWS the following information: descriptions of habitat type, growth form, maximum plant height and evolutionary (stem) age of the transition from the literature.

To obtain the number of DWS on continents and islands, we used the World Checklist of Vascular Plants (WCVP)^30,31^ for taxonomic standardization ^32^. We directly matched 6,295 species to WCVP using the taxize package v. 0.9.100 ^33^ package in R. We resolved an additional 425 species manually to WCVP (149 represented spelling variants and 276 had multiple matching names, differing in status and authority in WCVP and requesting a manual solution). A remaining 32 species from our dataset could not be matched with WCVP, and we disregarded them for subsequent analyses.

### The number of DWS and their geographic and phylogenetic distribution

To understand the geographic distribution of DWS, we used the information provided by the WCVP on the level of “botanical countries” (Level 3 of the World Geographic Scheme for Recording Plant Distributions)^34^. We chose this geographic scale since (1) the data are available for all vascular plants, allowing us to work with the proportion of DWS in the regional floras to assess island and continental DWS hotspots, and (2) the data contain information on the geographic status of species distributions in each region (native vs. introduced). We first obtained the number of DWS per botanical country, and then obtained the proportion of DWS of all native eudicot species occurring in this botanical country as a measure of the contribution of DWS to the total relevant diversity in each region. We used eudicots as a reference, since monocots cannot produce wood following our definition and early diverging flowering plant lineages do not comprize DWS. For all analyses, we excluded instances where species were considered extinct in a botanical country (2,701 instances), introduced (225,503), or their occurrence was doubtful according to the WCVP (2,950), retaining 1,739,296 occurrences of species in botanical countries.

To compare the prevalence of DWS on islands and continents, we considered “islands” as botanical countries that represent oceanic islands and continental fragments currently separated from the continent by water (Fig 1). To test if DW is more prevalent on continents than on islands, we used an Analysis of Variance to compare the proportion of DWS in continental and insular botanical countries, respectively.

**Figure 1.**
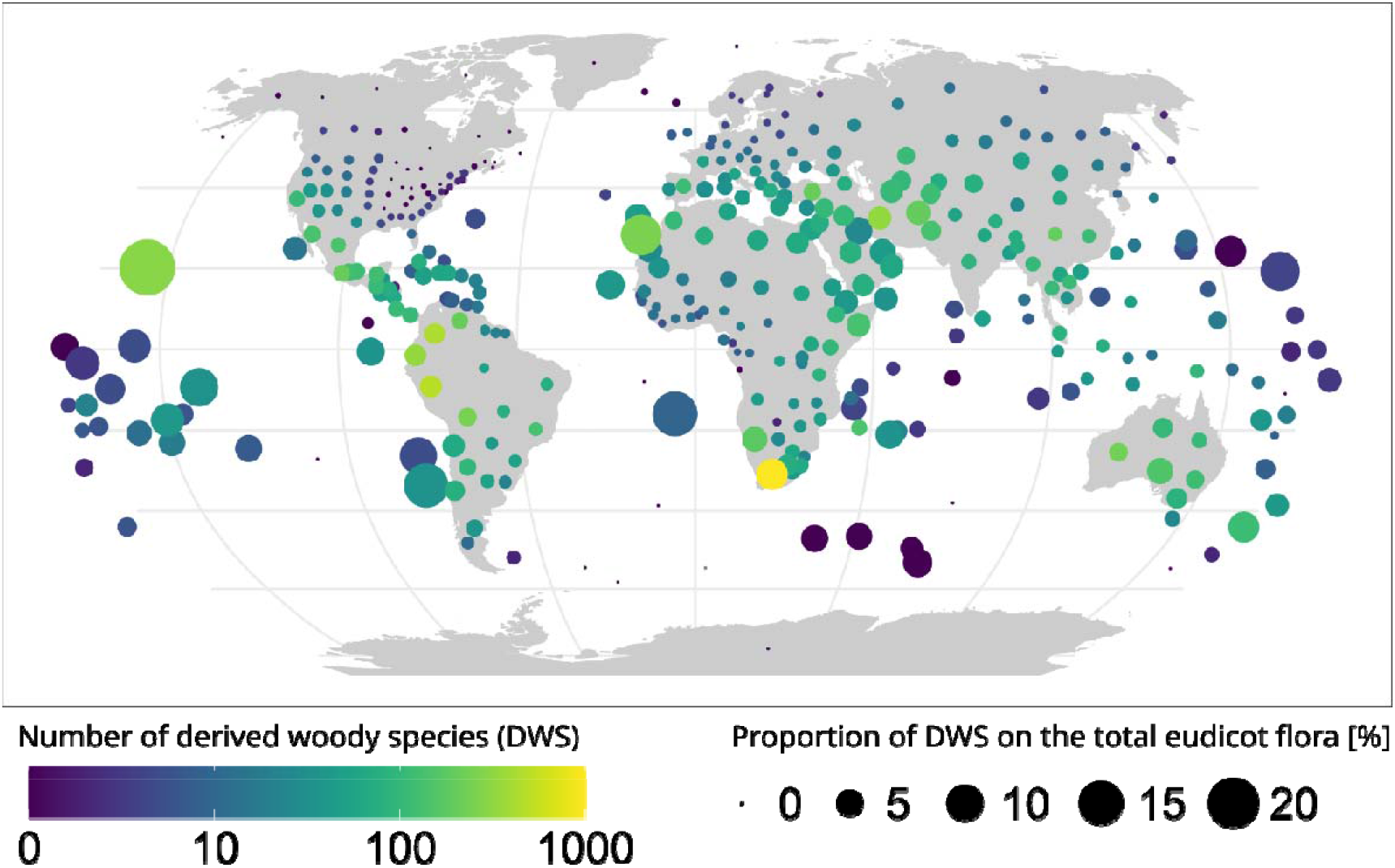
The global distribution of derived woody species (DWS) on islands and continents. Colour shows the absolute number of DWS in a region, the size of the bubble shows the proportion of DWS for the total native eudicot flora in that region. Islands are visualised only with a bubble; continental regions are plotted in the background.

To further understand the global distribution of DW, we searched for global centres of DWS richness with a null model approach. We used randomization of a species by botanical country matrix to generate 1,000 replicates on the number of expected DWS in a botanical country given its total native eudicot species richness, and then used a one-sided t-test at a significance level of 0.95 to identify botanical countries that have significantly more DWS than expected by their overall species richness. We identified DWS hotspots on continental and island areas separately.

To compare the phylogenetic distribution of DWS on islands and continents, we first summarized the number of DWS and the total number of species per genus and family. We then used the total number of species per taxon from WCVP to calculate the proportion of DWS, to identify the distribution of DW across the eudicot Tree of Life, and to test if specific clades are particularly prone to evolving DW. Furthermore, for those clades for which dated phylogenies were available, we obtained stem ages of derived woody lineages to compare clade ages, and grouped them for the two main IWS hotspots (Hawaii and Canary Islands) and the four continental DWS hotspots.

### Environmental correlates of DWS occurrence

To test the importance of drought and stable frost-free climate as predictors for DW, we related the proportion of DWS per botanical country to the summarized current environmental conditions per botanical country on continents and on islands. We obtained environmental data from the CHELSA v 1.2 data (only aridity index) ^35,36^ and the CHELSA v. 2.1 bioclim+ data (all other climate variables, aridity index not available in v2.1) ^37,38^ at 30 arc-second resolution and calculated their mean per botanical country, omitting raster cells with missing data. We selected mean aridity index (CHELSA acronym: ai, calculated as mean annual precipitation divided by mean annual evapotranspiration) and precipitation seasonality (bio15) to represent drought, and the number of growing degree days (ngd0, which are the days at which the mean daily air temperature is above 0°C) to represent climate related to frost. In addition to the climate variables, we obtained rasterized elevational data at 30 arc-seconds resolution from the Digital Elevation - Shuttle Radar Topography Mission ^39^, and added the botanical country’s mean elevation and the elevational range (i.e., the range from the 5 percentile to 95 percentile of all elevations recorded) as proxy for general habitat heterogeneity as environmental predictor. We used the distribution of recent species and current climate to test the links between DW and climate, assuming niche conservatism, since reliable paleoclimatic reconstructions of the relevant parameters for the time and location of the origin of DWS lineages were not available. To account for this, stem ages of derived woody lineages for six main DWS hotspots were mapped against major paleoclimatic events for these regions.

Some curation of the environmental data as obtained from CHELSA was necessary. Specifically, for the aridity index, we replaced raster-cell values smaller than 0.01 with 0.01 and values larger than 100 with 100. This affected only a small proportion of raster cells with unrealistically high values, due to the calculation of the aridity index as a quotient (meaning values get very large if potential evapotranspiration approaches zero, and very low if precipitation approaches zero). Furthermore, we imputed missing values for four instances for elevation and aridity index, respectively after summary on country level (for Howland-Baker Island, Marcus Island, Selvagens and St. Helena), using a random forest imputation as implemented in the missForest function of the missForest package in ^40^. These values were missing presumably due to the small size of the respective botanical countries, potentially combined with other factors limiting data retrieval, such as persistent cloud cover and isolated position in the ocean. We had also selected the annual range of air temperature (bio7) and the mean temperature of the growing season (gst) to additionally represent frost-free temperatures in the SEMs, but these variables were highly correlated with growing degree days (r > 0.7), and we therefore removed them from the analysis. Lastly, we log- or sqrt-transformed certain variables to improve normality and thereby model fit. Specifically, we sqrt-transformed the number of total eudicot species and log-transformed the number of derived woody species, the aridity index, the precipitation seasonality, the mean elevation and the total area, whereas we used the number of growing season days untransformed. We also normalized all variables between 0 and 1 to be able to compare the strength of effects driving DWS richness on continents compared to IWS richness on islands.

We then used structural equation models (SEMs) in the R package lavaan version 0.6.19 ^41^ to test for the drivers of the number of DWS species per region. We fitted three SEMs: one including all continental and island botanical countries (N = 368) to estimate the overall effect of the environmental variables, one exclusively for continental botanical countries to estimate the effect of the environmental variables on continents (N = 244), and one including only islands (N = 124). Since we were interested in the environmental factors driving the presence of DWS, we included all DWS of a botanical country even if they putatively evolved their woodiness elsewhere. For example, we included DWS that occur on islands today, but evolved DW on a nearby continent. We started with an a priori SEM that included all hypothesized pathways among predictor variables. Specifically, we assessed how the selected environmental variables and total area of the region affected total eudicot species richness, and how the selected variables and total plant richness affected variation in the number of DWS on continents and IWS on islands. In other words, we tested the direct effects of the environmental variables on the number of DWS and IWS, as well as the indirect effects of these variables via their effect on total species richness. We progressively deleted paths with the least statistical significance from the SEMs until our final model only consisted of significant pathways (at p < 0.05), for which we extracted the standardized coefficients. For this final model, we evaluated model’s modification indices, model fits, and residual correlations ^42^ To ensure adequate fit of the SEMs, we evaluated the P values of χ2 tests, the comparative fit index and confidence intervals of the root mean square error.

### Habitat, maximum plant height, and growth form

To assess whether the occurrence of DWS is related to more open habitats, we obtained information on habitat (available for 3,870 DWS) and species elevational range (available for 3,114 DWS) from the literature when possible, and converted standardized free text descriptions into broader habitat types (Table 1) based on key words. Additionally, we obtained maximum plant height (available for 4,390 species) and growth form (available for 6,507 species) per species from the literature when available, and used Analyses of Variance to test if DWS’ plant height is different between islands and continents.

**Table 1.**
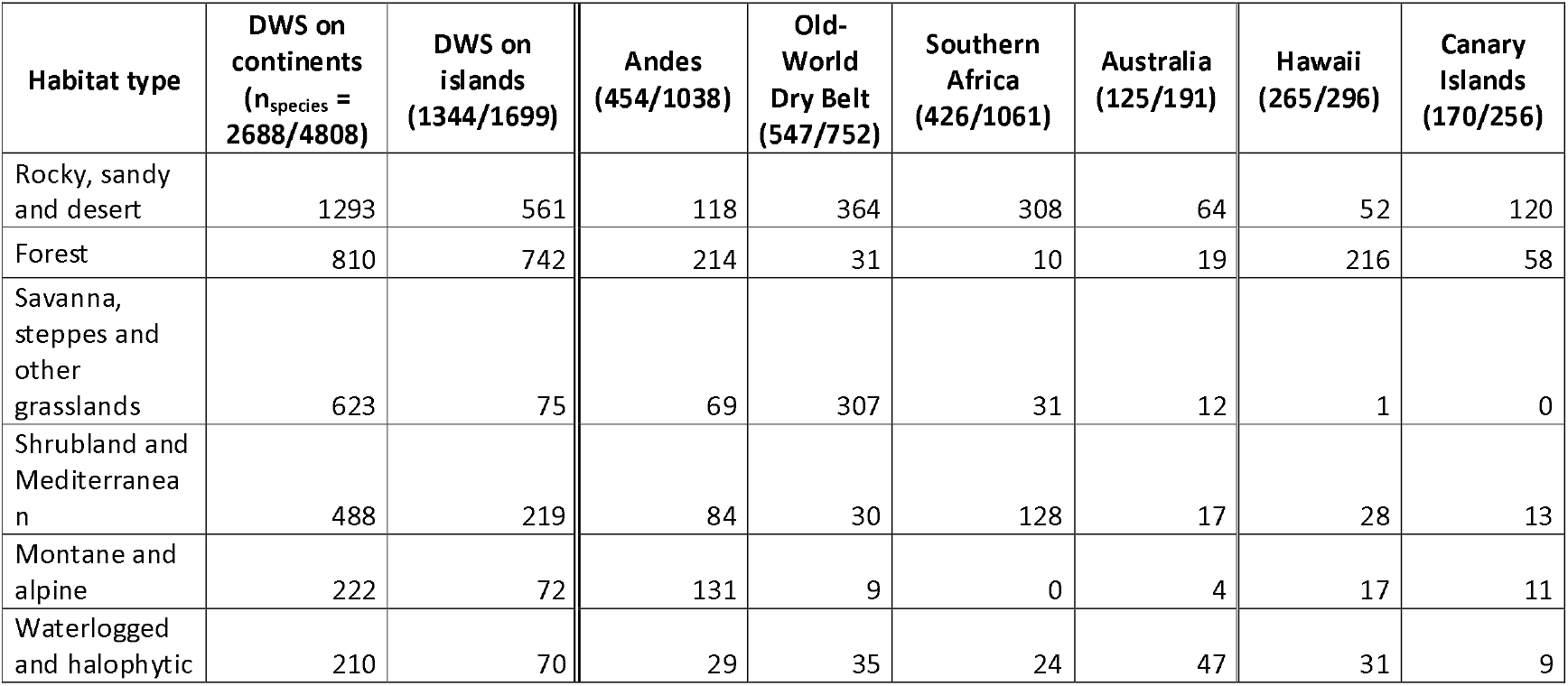
Habitat preference of derived woody species (DWS) on islands and continents and in the main continental and insular DWS hotspots, based on 3,870 DWS for which habitat information was available (species numbers per total DWS number provided in each of the categories). Species may occur in multiple habitat types.

| Habitat type | DWS on continents<br>($n_{\text{species}} = 2688/4808$) | DWS on islands<br>(1344/1699) | Andes<br>(454/1038) | Old-World<br>Dry Belt<br>(547/752) | Southern<br>Africa<br>(426/1061) | Australia<br>(125/191) | Hawaii<br>(265/296) | Canary<br>Islands<br>(170/256) |
| --- | --- | --- | --- | --- | --- | --- | --- | --- |
| Rocky, sandy and desert | 1293 | 561 | 118 | 364 | 308 | 64 | 52 | 120 |
| Forest | 810 | 742 | 214 | 31 | 10 | 19 | 216 | 58 |
| Savanna, steppes and other grasslands | 623 | 75 | 69 | 307 | 31 | 12 | 1 | 0 |
| Shrubland and Mediterranean | 488 | 219 | 84 | 30 | 128 | 17 | 28 | 13 |
| Montane and alpine | 222 | 72 | 131 | 9 | 0 | 4 | 17 | 11 |
| Waterlogged and halophytic | 210 | 70 | 29 | 35 | 24 | 47 | 31 | 9 |

### Software used

We used R version 4.4.2 ^43^ and the following R packages: adephylo v. 1.1.16, ape v. 5.8.1, car v. 3.1.3, cowplot v. 1.1.3, deeptime v. 2.1.0, exactextractr v. 0.10.0, ggnewscale v. 0.5.0, ggtree v. 3.14.0, gridExtra v. 2.3, janitor v. 2.2.1, measurements v. 1.5.1, missForest v. 1.5, phytools v. 2.4.4, raster v. 3.6.31, RColorBrewer v. 1.1.3, rmapshaper v. 0.5.0^44^, rnaturalearth v. 1.0.1, rWCVP v. 1.2.4, WCVPdata v. 0.5.0, scales v. 1.3.0, sf v. 1.0.19, sp v. 2.2.0, taxize v. 0.10.0, terra v. 1.8.21, tidyverse v. 2.0.0, viridis v. 0.6.5, wesanderson v. 0.3.7, writexl v. 1.5.1. See Supplementary Material S1 for the software references.

## Results

### The number of DWS and their geographic and phylogenetic distribution

We found a total of 6,507 DWS in 647 genera and 46 families. Of these, 4,808 evolved woodiness on continents (74%) with at least 513 evolutionary transitions, versus 1,084 on islands (IWS, 17%) with at least 175 transitions. Another 615 DWS were endemic to islands, but their lineage likely evolved woodiness on nearby continents before establishing on the island (9%).

DWS occurred in 337 out of 369 botanical countries, with virtually no DWS in high-latitude regions. On continents, the botanical countries with most DWS were the Cape Provinces of South Africa (994 species), Peru (483), and Colombia (433), and the proportion of DWS was highest in the Cape Provinces of South Africa (6.86%), Kuwait (5.07%) and South Australia (4.59%; Fig. 1). On islands, the number of DWS (including IWS and continental DWS that later dispersed to islands) was highest on the Hawaiian archipelago (296), Canary Islands (256), and Madagascar (144), while the proportion of DWS was highest on the Hawaii archipelago (24.2%), St. Helena (15.3%), and the Juan Fernández Islands (14.8%). The mean total number of DWS was higher in the larger continental botanical countries (mean = 52.3) than in the smaller islands (mean = 22.5, F_1,366_[12.3], p < 0.001). However, the proportion of DWS was on average higher on islands (mean = 2.73% vs. 1.02% in continental regions; F_1,366_[48.5], p < 0.0001). We identified four global continental DWS hotspots composed of multiple adjacent botanical countries with significantly higher DWS proportions than expected (Fig. 2): Southern Africa (1,061 DWS/6.35%/at least 152 independent evolutionary transitions), the Andes (1038/3.33%/188), the Old-World Dry Belt (752/3.31%/206), and Australia (191/2.56%/58). For islands, we identified 22 DWS hotspots, with the Hawaiian archipelago (296/24.2%/17) and the Canary Islands (256/12.1%/33) standing out due to their high absolute number of DWS and evolutionary transitions.

**Figure 2.**
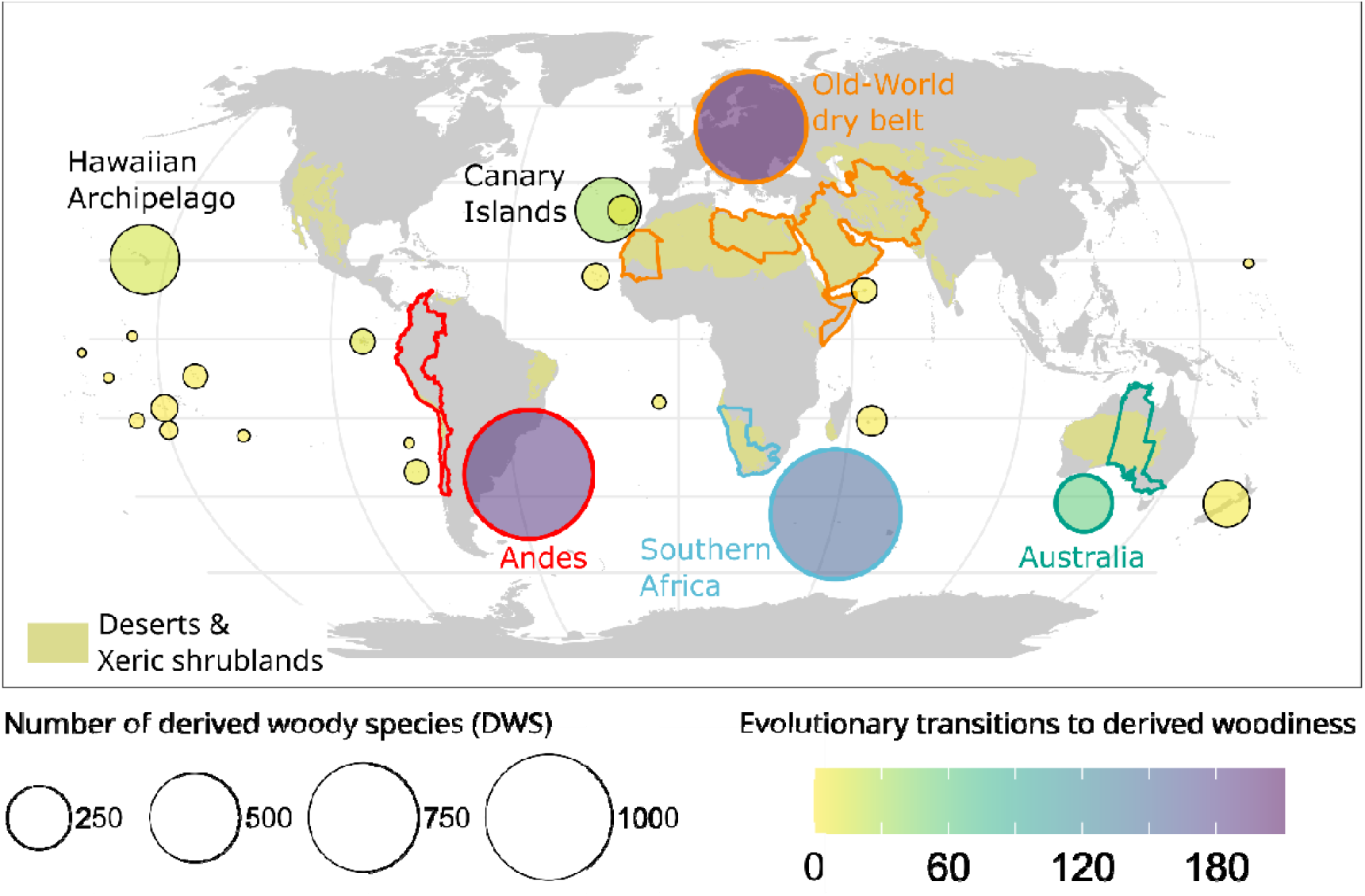
Global centres of derived woodiness (DW) across the globe, showing absolute numbers of derived woody species (DWS) and evolutionary transitions towards DW. Based on a randomization test, botanical countries in four continental regions and 22 archipelagos emerged as global DWS hotspots where the number of DWS is significantly higher than expected given total native eudicot species richness. Concerning islands, the Hawaiian archipelago and Canary Islands stand out with a high absolute number in DWS and evolutionary shifts. The continental DWS hotspots overlap considerably with the desert and xeric biome.

DWS were absent in the earlier diverging eudicot lineages, but scattered across the remaining eudicot Tree of Life, with clustering in the super-asterids (Fig. 3). Most families were of similar importance on continents and islands. Asteraceae were outstanding, with a high number of DWS on continents (973) and islands (388), as well as the highest number of minimum transitions on continents (94) and islands (47). The remaining top families with DWS were Brassicaceae, Aizoaceae, Amaranthaceae and Lamiaceae (Fig. 3). Notably, the number of DWS in the individual families varied among the different geographic DWS hotspots (Fig. 3).

**Figure 3.**
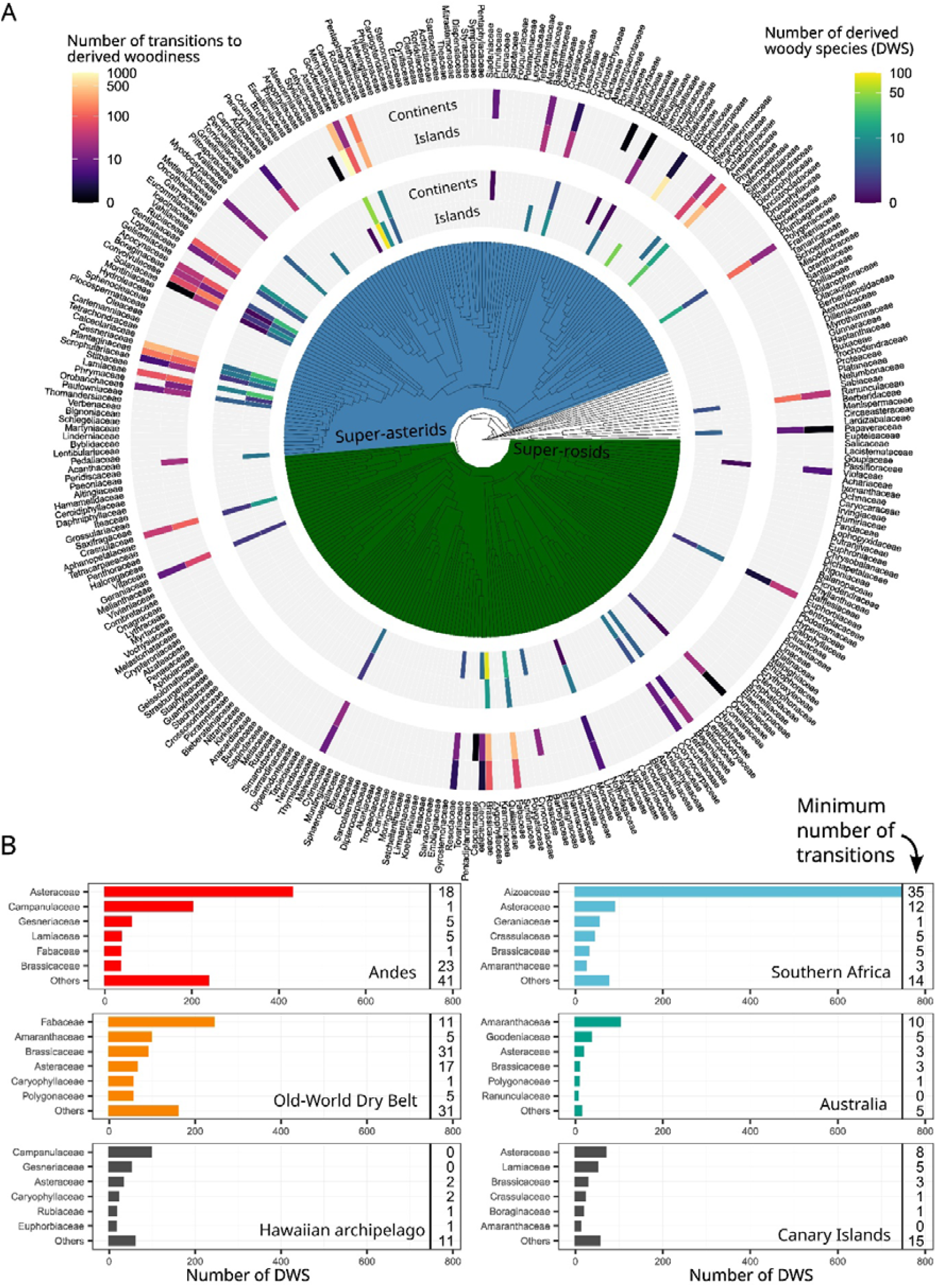
Distribution of derived woody species (DWS) on the eudicot Tree of Life^58^. **A)** The number of DWS and evolutionary transitions to DW per family in the phylogenetic context. **B)** The number of DWS and minimum number of evolutionary transitions in the main continental and insular DWS hotspots for each of the six families with the most DWS.

Most DW lineages originated in the Plio-Pleistocene, and stem ages of DW lineages ranged from 48 (*Aethionema*, Brassicaceae and *Oxalis*, Oxalidaceae) to 0.1 (*Lotus*, Fabaceae) million years ago, with an average age of 7.74 Ma (Fig. 4). On average, insular woody lineages were younger than continental derived woody lineages (average stem age 5.18 Ma vs. 9.36Ma; F_[192, 1]_ = 12.2, p<0.001).

**Figure 4.**
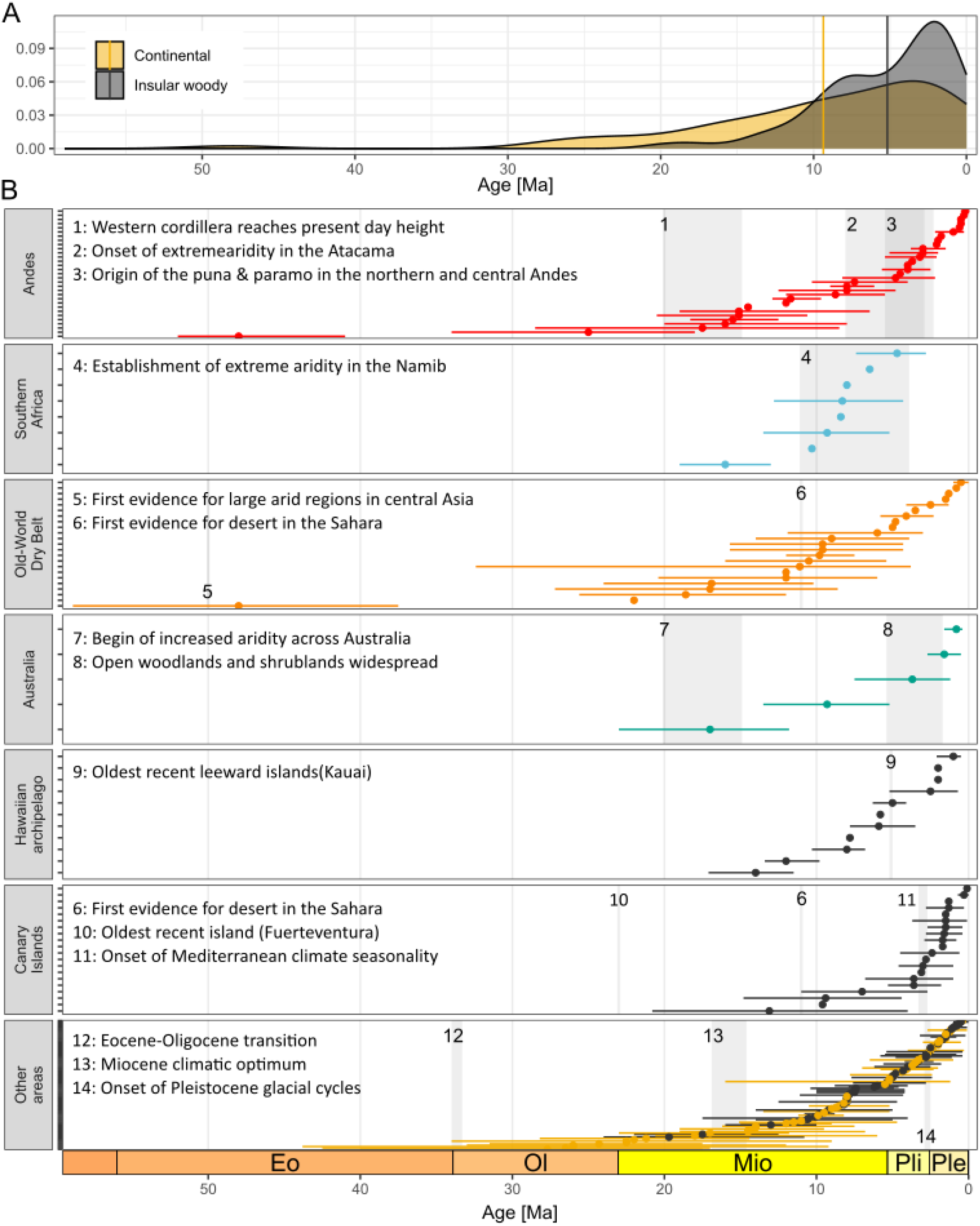
The (stem) age of derived woody clades (N_lineages_ = 192). **A)** Average age of insular woody and continentally woody clades. **B)** Average age across major derived woodiness hotspots. The grey bars and numbers indicate the estimated minimum ages of selected regional paleoclimatic and geological events. See Supplemetary Material S3 for the references.

### Environmental predictors of DWS distribution on islands and continents

Several variables were significant predictors of the occurrence of DWS on continents and/or islands (Fig. 5). In all cases, the total eudicot species richness was the most important predictor of DWS richness (Fig. 5A), along with precipitation seasonality, number of growing season days and mean elevation. Hence, the number of DWS is on average higher in botanical countries with higher overall eudicot richness, higher precipitation seasonality, more growing season days and higher elevation, irrespective of island or continental setting. In contrast, the aridity index was a strong negative direct predictor for the number of DWS only on continents (but not significant for islands), and the number of growing season days had a larger effect on islands (Fig. 5A).

**Figure 5.**
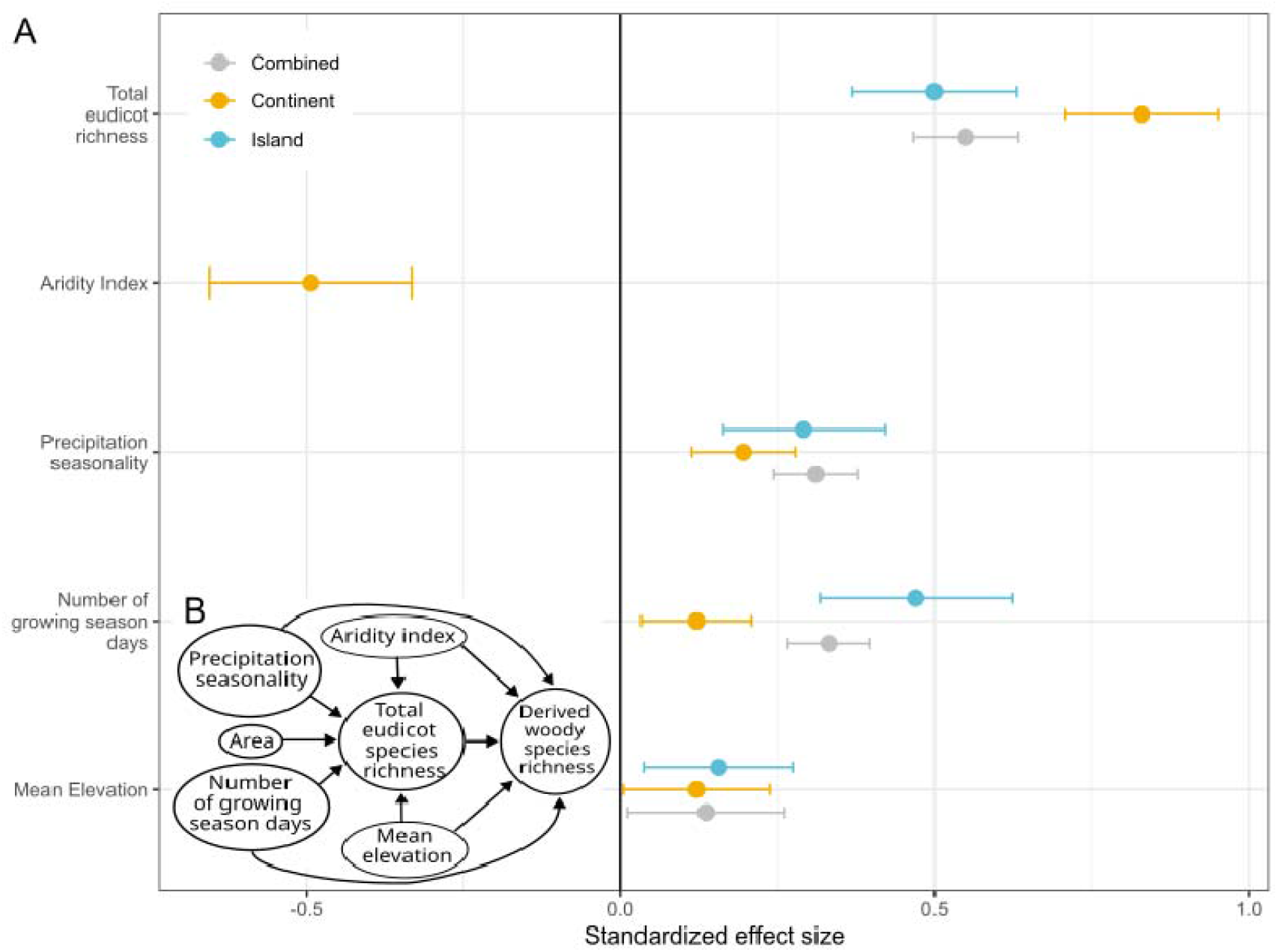
Results of three structural equation models (SEMs) relating the number of derived woody species (DWS) to average environmental conditions across botanical countries worldwide. **A)** Standardized model coefficients of three individual models including all botanical countries (grey, n = 368), continental botanical countries (orange, n = 244) and island botanical countries (blue, n = 124). Only significant coefficients of the direct effects are shown. **B)** Model structure for the initial models including all effects. For the combined model the direct effect of aridity on the number of DWS was not significant, and for the island model neither the indirect effect of ‘Mean elevation’ nor any effects of the ‘Aridity index’ were significant.

The combined, continental and island models explained 64/71/66% variation in DWS richness, respectively. The environmental variables also explained considerable variation in total eudicot species richness (69/57/80%, respectively). Overall model fit was good for the continent and island model with P values of χ2 tests greater than 0.05 (continent model: 0.35; island model: 0.62), with comparative fit index > 0.90 (continent model: 1.0; island model: 1.0) and confidence intervals of the root-mean-square error of approximation (continent model: lower: 0.00, upper: 0.17; island model: lower: 0.00, upper: 0.143). Fit was less good for the combined model with P values of χ2 tests < 0.05, comparative fit index at 0.98 and confidence intervals of the root-mean-square error of approximation at 0.08 (lower) and 0.186 (upper).

### Distribution of DWS in open-closed vegetation types, maximum plant height and growth form

On continents and islands, DWS occurred more often in open habitats than in forests, comprising 87% and 57% of the species for which habitat data were available, respectively (Table 1). However, the number of DWS in different habitat types varied among the DWS hotspots, with most DWS in Australia, the Old-World Dry Belt, Southern Africa and the Canary Islands occurring in “rocky, sandy or desert” habitats and most DWS in the Andes and the Hawaiian archipelago occurring in “forest”. Notably, of the DWS on continents, 1,929 occurred above 1000m elevation irrespective of habitat type, and 1,484 exclusively above 1000m (Fig. 6). Elevational preferences also differed among geographic DWS hotspots, with the DWS in the Andes preferring higher elevations in alpine or montane habitats, in congruence with the increased availability of these habitats compared to other regions (Fig. 6A).

**Figure 6.**
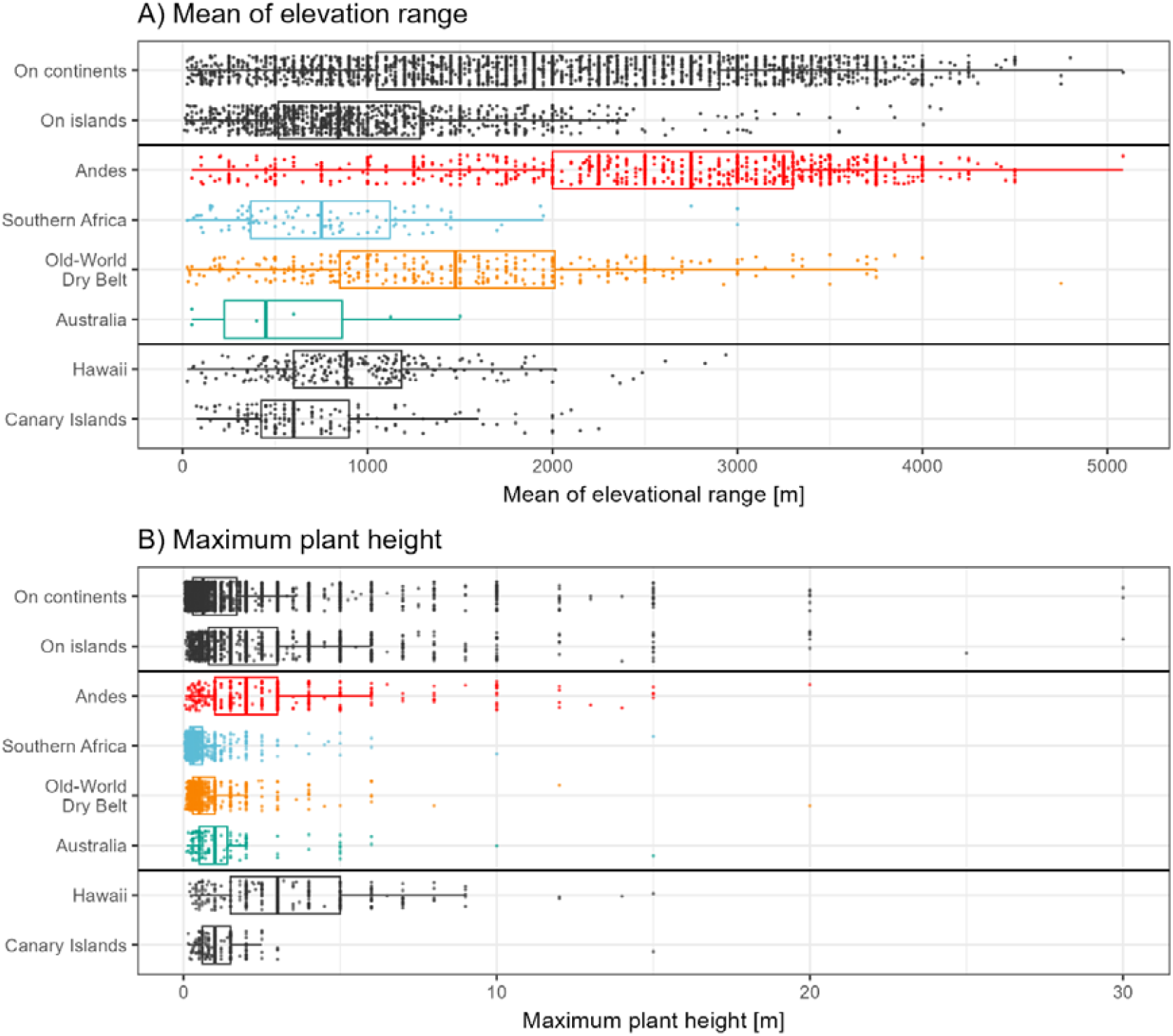
Characteristics of derived woody species on continents and islands and in the six global DWS hotspots. **A)** Mean of the elevation range, including 3,114 species for which data was available. **B)** Maximum plant height based on information from 4,390 DWS.

DWS on islands grew on average taller than on continents (mean 2.3 m and 1.6 m, respectively; F_[4288, 1]_ = 52.6, p < 0.0001, Fig. 6B). Also plant height differed significantly among continental DWS hotspots (F_[2760, 1]_ = 48.61, p < 0.0001, Fig. 6B): DWS on average were tallest on the Hawaiian archipelago (3.3 m) and in the Andes (2.8 m), and lower in Australia (1.3 m), Canary Islands (1.2 m), Old-World Dry belt (0.9 m), and Southern Africa (0.6 m). At the global scale, roughly 85% of DWS were shrubs, with most remaining species on islands being trees or treelets (8% and 5% respectively), and climbers on continents (10%). Notably, the proportion of climbers was elevated in the Andes where 19% were climbers.

## Discussion

The rampant evolutionary transitions towards woodiness reveal that derived woody species (DWS) are much more abundant and widespread outside island archipelagos than previously thought (Figs. 1-3, Table 1) ^4,13^. Our woodiness database reveals that the number of continental DWS is almost five times higher than suggested by previous scattered literature reports ^14–22^, indicating that continents harbour 83% of the total 6,500 DWS and 75% of the nearly 700 transitions, most of them in the Andes, Southern Africa and the Old-World Dry Belt (45% of the total number of DWS) plus 3% in the fourth continental hotspot represented by the arid-Mediterranean regions of Australia. Notably, the proportion of DWS on the total regional eudicot flora is often higher on islands, resulting in 22 archipelago hotspots, most prominently the Hawaiian archipelago and the Canary Islands (Fig. 2). The actual numbers of DWS and transitions to woodiness are likely even higher, given our conservative definition of woodiness and the lack of densely sampled, robust phylogenies for many angiosperm clades.

Several families – especially in the superasterids clades (e.g., Asteraceae, Aizoaceae, Amaranthaceae) and in Brassicaceae – seem to be more likely to undergo woodiness transitions, and repeatedly evolved DWS in various parts of the world at different times (Figs. 3-4). Similarly, at the genus level, prime examples for multiple transitions are *Salvia* (Lamiaceae, 19 transitions leading to 68 DWS mainly in the Andes above 2000m, in South Africa and some parts of the Old-World dry belt, and in Canary Islands and Madagascar), and *Lepidium* (Brassicaceae, 7/45 on various islands in the Pacific, and in dry parts of all four continental DWS hotspots as well as southwestern USA). In other genera with multiple DW transitions, transitions are more confined to individual DWS hotspots such as in *Ptilotus* (Amaranthaceae, 7/26 in interior deserts of western Australia) and *Valeriana* (Caprifoliaceae, 5/27 in paramo of the Andes), or limited to archipelagos like in *Euphorbia* (Euphorbiaceae, 7/37 in Macaronesia, Hawaii and Galapagos Islands) and Sonchus (Asteraceae, 3/32 in Macaronesia, Juan Fernandez Islands, San Ambrosio Islands). There are also examples of single woodiness transitions that radiated into multiple DW genera, such as the Espeletia clade in the Andes (Asteraceae, 7 genera including 119 DWS) ^45^, the *Sclerolaena* clade in Australia (Amaranthaceae, 7 genera leading to 73 DWS)^46^, and the iconic lobelioid clade native to the Hawaiian archipelago, the southern Pacific and the African mountains (Campanulaceae, 8 genera resulting in 160 DWS) ^17,47^. Interestingly, in many of these large DW radiations, multiple reversals from DW back to secondarily derived herbaceousness have happened.

Overall, there is no obvious connection between woodiness transitions and species richness. Despite the above-described examples of species-rich derived woody lineages and the proposed role of DW as a key innovation that increases diversification in selected DW lineages ^48^, woodiness transitions have not necessarily resulted in radiations across lineages. Indeed, 37% (257 out of 866) of the lineages that evolved DW only include one or two DWS. Interestingly, these 257 lineages proportionally overrepresent island clades compared to continental ones (50% and 32%, respectively). This may be linked to higher extinction rates in IW lineages due to the smaller and geologically more dynamic islands, and/or older average age in the continental DW lineages leaving more time for species accumulation (Fig. 4A). Consequently, traits other than DW are likely involved in the diversification dynamics of DW lineages.

Drought is the most important global woodiness driver on continents. Three of the four global continental DWS hotspots (southern Africa, Old-World dry belt, Australia) are characterized by long and intense drought cycles, and largely comprise of deserts, steppes and xeric shrublands (Fig. 2). Interestingly, most of the dated DW lineages in these three DWS hotspots might have originated in paleoclimatic periods associated with increased drought (Fig. 4B). Likewise, the bulk of DWS in southern Africa thrives in the western winter-rainfall area with more prolonged and intense drought periods compared to the eastern area ^49^ The drought hypothesis is also supported by experimental studies that found a positive correlation between increased stem woodiness and resistance to lethal levels of drought-induced gas embolism in the vasculature in multiple lineages native to islands and continents ^12,50^. Our SEM results confirm a strong correlation between the occurrence of DWS and seasonal drought on continents, also when accounting for total species richness (Fig. 5). However, drought is not the sole woodiness driver as the global DWS distribution pattern is more complex at finer taxonomic and regional scales. For instance, the Andes ecosystem, which is the most species-rich global continental DWS hotspot, exhibits the most variation in terms of environmental variables, suggesting additional drivers have probably played significant roles in woodiness evolution on these tropical mountain peaks (see below for a more elaborate discussion). In summary, drought has been a major global driver of woodiness evolution, especially on continents, but more detailed results will reveal multiple lineage-specific drivers to account for each of the 700 evolutionary transitions.

Frost-free temperatures are the main driver of DWS on islands, while playing a secondary role on continents. The positive impact of frost-free climate is evident based on the decline of the number of DWS species towards higher-latitude regions on continents and islands ^13^. This is supported by the SEM showing a positive correlation between the number of growing season days (proxy for frost-free period) and the occurrence of DWS, especially on islands and to a lesser extent also on continents (Fig. 5). The number of frost days also emerged as a predictor of the number of IWS on the more isolated oceanic islands in a previous study using different data on plant occurrences ^13^. The positive link between frost-free temperatures and DW relates to the favourable aseasonal climate hypothesis postulated for IW ^4^. In addition, already more than 100 years ago, Sinnott and Bailey ^51^ argued that many woody angiosperm lineages evolved into herbaceous lineages due to declining temperatures during the Paleogene and Neogene that gave rise to cold winters in the temperate regions. Likewise, in extant angiosperms, freezing environments favour either herbaceous growth or a few woody species with narrow vessels and/or deciduous leaves ^52^. The only DWS that experience frequent frost events are confined to tropical alpine habitats typically above 3400-3900m asl (at least 300 DWS, see Table 1; Fig. 6A), and exhibit several morphological and physiological frost-associated adaptations ^53^. Interestingly, the development of tall aerial stems in rosette trees is thought to represent an adaptation that protects meristematic and reproductive tissues from the colder temperatures occurring at ground level compared to higher above the surface. Several other potential drivers potentially play a role in woodiness evolution on these continental sky islands as well (see next section). Overall, increasing evidence indicates that frost constrains the evolution of woodiness globally, with a particularly strong impact on islands.

The abundance of DWS in continental mountain systems suggests various potential drivers that stimulate woodiness evolution. High-elevation areas in the (sub)tropical belt are often rich in DWS, with the Andes as a prime example (Table 1; Figs. 5; 6A). However, the variable we used in the SEM (mean elevation) is a proxy for many parameters that change with elevation, which complicates our understanding of the mechanisms involved in woodiness evolution on tall tropical mountains. Indeed, the Andes are the most variable DWS hotspot in terms of environmental conditions, based on the precipitation difference between the drier western flanks and the wetter eastern flanks, and the steep altitudinal range with several hundreds of DWS growing at alpine elevation above 3500m asl (Fig. 5A). Among the various potential woodiness drivers that are involved along the flanks of the Andes, drought could be one of them. At least 25% of the DWS in the Andes occur in habitats with a clear drought signal (rocky, sandy, desert, savanna or grasslands, shrubland or Mediterranean; Table 1). Additionally, available literature confirms that pollinator diversity in the Andes decreases with elevation ^54^, which links to one of the IW hypotheses stating that woodiness is stimulated in pollinator-poor regions ^4^, although this hypothesis remains to be experimentally tested. Another aspect that requires more investigation is how mutagenic effects from solar radiation in the open tropical alpine environments impacted woodiness. Speculatively, perennial species with woody stems surrounded by bark contain more phenylpropanoid-derived substances, including lignins and UV-B absorbing flavonoids, that better protect them against ultraviolet radiation compared to shorter-lived herbs ^55^. In summary, multiple global woodiness drivers are likely at play, especially in environmentally variable DWS hotspots, but more experimental work is required to tease apart the impact of these potential drivers in promoting woodiness evolution.

The majority of DWS thrive in open habitats, suggesting woodiness evolution may also be impacted by competition among plants, as already hypothesised by Darwin ^5^. This is particularly evident in the diverse continental lowland rainforests where DW is virtually absent (Fig. 2), likely because tall ancestrally woody trees outcompete the much smaller DWS for light, which in turn may prevent woodiness evolution in the dark understorey ^16^. This does not mean, however, that DWS are absent in forests. Especially islands harbour a substantial proportion of DWS in forests (on average 45% on islands vs. 17% on continents), although there are large differences among archipelagos (75% of all DWS on the Hawaiian archipelago occur in forests vs. 23% on the Canary Islands) and across continental DWS hotspots (20% in Andes vs. 10/4/1% for Australia, Old-World Dry Belt and South

Africa, respectively; Table 1). Overall, more than half of the DWS on continents and islands thrive in open vegetation types (e.g., deserts, steppes, savannas, salt planes, and Mediterranean scrubland), which are most dominant in the continental DWS hotspots of the Old-World Dry Belt (98%), Australia (75%), and southern Africa (more than 50%; Table 1). Whether or not these light-stimulating open environments are simply the result of the strong seasonal drought, and/or rather promote woodiness to grow into taller shrubs, is hard to disentangle. Interestingly, DWS are on average shorter on continents compared to islands (1.6 vs. 2.3 m, respectively; Fig. 6B), probably due to the stronger association of continental DWS with drought. Although this woody growth habit remains relatively short in stature, it is noteworthy that DWS on continents — and particularly on islands — are likely considerably taller than co-occurring herbaceous species, given that most herbaceous species worldwide are short ^56,57^.

In conclusion, our global review on the biogeography and evolution of phylogenetically derived woodiness in angiosperms reveals that DW is common across islands and continents, with a minimum number of almost 700 independent transitions, particularly across 22 (sub)tropical archipelagos and four continental DWS hotspots. Our results suggest that, at a global scale, drought is probably the most important driver of derived woodiness on continents, whereas frost-free temperatures have a higher impact on islands. In addition, the biological role of co-occurring species competing for light may have favoured woodiness transitions, as suggested by the considerable height of many of the shrubby DWS – especially on islands – and their predominant occurrence in open habitats. Hence, DW can be shaped by multiple factors, and we anticipate that future experimental studies examining DWS at finer taxonomic and spatial scales will uncover a more complex array of lineage- and region-specific woodiness drivers. In addition to (a)biotic drivers involved in DW across the various DWS hotspot areas, intrinsic aspects of the herbaceous lineages (e.g., pre-adaptation of lineages, genomic rearrangements) are important to determine whether or not a given clade develops DW and/or undergoes radiation.

## Supporting information

Supplementary file 1 - Additional references

## Data availability

All data and scripts used for the analyses presented in this study will be available at publication.

## Acknowledgements

M.G. acknowledges the support of the German Centre for Integrative Biodiversity Research (iDiv) Halle - Jena - Leipzig, funded by the German Research Foundation DFG–FZT 118, grant 202548816.

## Author contributions

AZ, FL, RO, HB designed this research. FL collected the data. AZ, MG and REO analysed the data, with contributions from TC. AZ and FL wrote the manuscript with contributions from all authors.

