## Supplementary file 1 - Additional references for "Global convergence in wood evolution is driven by drought on continents and frost-free temperatures on islands"

Supplementary material 1, Zizka et al. (2025) - Additional references

### Software used

We used R version 4.4.2 (R Core Team, 2024) and the following R packages: adephylo v. 1.1.16 (Jombart & Dray, 2010), ape v. 5.8.1 (Paradis & Schliep, 2019), car v. 3.1.3 (Fox & Weisberg, 2019), cowplot v. 1.1.3 (Wilke, 2024), deeptime v. 2.1.0 (Gearty, 2024), exactextractr v. 0.10.0 (Daniel Baston, 2023), ggnewscale v. 0.5.0 (Campitelli, 2024), ggtree v. 3.14.0 (Yu et al., 2017), gridExtra v. 2.3 (Auguie, 2017), janitor v. 2.2.1 (Firke, 2024), measurements v. 1.5.1 (Birk, 2023), missForest v. 1.5 (Stekhoven & Buehlmann, 2012), phytools v. 2.4.4 (Revell, 2012), raster v. 3.6.31 (Hijmans, 2025a), RColorBrewer v. 1.1.3 (Neuwirth, 2022), rmapshaper v. 0.5.0 (Teucher & Russell, 2023), rnaturalearth v. 1.0.1 (Massicotte & South, 2023), rWCVP v. 1.2.4 (Brown et al., 2023), WCVPdata v. 0.5.0 (Govaerts, 2024), scales v. 1.3.0 (Wickham et al., 2023), sf v. 1.0.19 (Pebesma & Bivand, 2023), sp v. 2.2.0 (Bivand et al., 2013), taxize v. 0.10.0 (Scott Chamberlain & Eduard Szocs, 2013), terra v. 1.8.21 (Hijmans, 2025b), tidyverse v. 2.0.0 (Wickham et al., 2019), viridis v. 0.6.5 (Garnier et al., 2024), wesanderson v. 0.3.7 (Ram & Wickham, 2023), writexl v. 1.5.1 (Ooms, 2024).

### Timing of paleoclimatic events

The grey bars and numbers indicate the estimated minimum ages of selected regional paleoclimatic and geological events: 1: Western cordillera reaches present day height (Kiel et al., 2023; Pérez-Escobar et al., 2022), 2: Onset of extreme aridity in the Atacama region (Garreaud et al., 2010), 3: Origin of the puna and paramo alpine ecosystems in the Northern and central Andes (Martínez et al., 2020), 4: Establishment of extreme aridity in the Namib region (Huntley, 2023), 5: First evidence of large arid regions in central Asia (Zhang et al., 2022), 6: First evidence for desert in the Sahara region (Crocker et al., 2022), 7: Beginning of increased aridity across Australia (Fujioka & Chappell, 2010), 8: Open woodlands and shrublands widespread (Fujioka & Chappell, 2010), 9: Oldest leeward recent island (Kauai) (Blay & Chuck, 2004), 10: Oldest recent island (Fuerteventura) (van den Bogaard, 2013), 11: Onset of Mediterranean climate seasonality (Suc, 1984), 12: Eocene-Oligocene Transition, 13: Miocene climatic optimum, 14: Onset of Pleistocene glacial cycles.
